# Acoustic niche partitioning in two tropical wet forest bird communities

**DOI:** 10.1101/2020.08.17.253674

**Authors:** Patrick J. Hart, Kristina Paxton, Thomas Ibanez, Grace Tredinnick, Esther Sebastián-González, Ann Tanimoto-Johnson

## Abstract

When acoustic signals sent from individuals overlap in frequency and time, acoustic interference and signal masking may occur. Under the acoustic niche hypothesis (ANH), signaling behavior has evolved to minimize overlap with other calling individuals through selection on signal structure and the sender’s ability to adjust the timing of signals. In this study, we examine the fine-scale use of acoustic space and the relevance of the acoustic niche hypothesis in two montane tropical wet forest bird communities (Costa Rica and Hawai’i) that vary in bird species richness. We used a null model approach to test the prediction that there are differences between observed and expected signal overlap in both communities. As predicted under ANH, we found much lower overlap of acoustic signals than expected by chance. In addition, spectral and temporal overlap between different signals was far more common in Hawaii than Costa Rica. These findings constitute strong support that there is competition for acoustic space in signaling communities, and this has resulted in temporal and spectral partitioning of the soundscape.

## Introduction

Acoustic signaling is a major form of social behavior in many terrestrial and aquatic organisms. When acoustic signals sent from individuals overlap in frequency and time, acoustic interference and signal masking occurs, which may reduce the receiver’s ability to discriminate information from the signal [1, 2]. Under the acoustic niche hypothesis (ANH; 3, 4), signaling behavior has evolved to minimize overlap with heterospecific calling individuals through selection on signal structure and the sender’s ability to adjust the timing of signals. This hypothesis may be viewed as an extension of the niche theory of Hutchinson [5] whereby acoustic space is a resource that organisms may compete for and that can be partitioned both spectrally (frequency range of the signal) and temporally.

The ANH is central to describing how animal signals in diverse calling communities are dispersed in space and time and is a major organizing hypothesis in the field of soundscape ecology [6]. While the ANH makes intuitive sense and is based on both anecdotal and empirical evidence from a broad range of studies and signaling taxa [7], the degree to which organisms partition acoustic space in order to reduce interference remains the subject of recent debate [8]. Studies in biodiverse calling communities such as cicadas [9], crickets [10], anurans [11-13], and birds [14-17] have found evidence for niche partitioning through the apparent evolution of signal character displacement among species. Other studies have demonstrated that birds can adjust the fine-scale timing of their signals to take advantage of gaps in noise and thus minimize temporal acoustic overlap with other birds [18-21] and even insects [22]. In all cases however, support for ANH can only be considered partial. Those studies that have found evidence of spectral character displacement of the signals themselves were not able to determine if that resulted in a significant reduction in overlap with other signals in real time. Those studies that reported temporal adjustments in signals generally did so for select species groups and not the signaling community as a whole. In order to have the clearest test of ANH, it is important to determine how organisms simultaneously partition both the frequency and the timing of their vocalizations in real time and at the community level.

Similar to the way other limiting resources (and thus niche partitioning) vary across space and time, acoustic niche space would be expected to be partitioned in some habitats, communities, and time periods, but not in others. Tropical wet forests have the most species-rich assemblages of organisms that signal acoustically, and thus competition for acoustic niche-space is expected to be strongest there [14, 23]. Conversely, acoustic space is not expected to be limiting in ecosystems that have a low richness or density of signaling organisms. In this study, we examine the fine-scale use of acoustic space and the relevance of the acoustic niche hypothesis in montane tropical wet forest bird communities in both Costa Rica and Hawaii. We used a null model approach to test the major prediction under the ANH that there are differences between observed and expected levels of signal overlap in both communities. We also examined the way the use of acoustic space differs between the two bird communities, and predicted that the degree of partitioning is greater in the species-rich community due to selection to reduce inter-specific signal overlap.

## Materials and methods

### Study sites

The Costa Rica study site was within an approximately 360 ha forest fragment at the Organization for Tropical Studies (OTS) Las Cruces Biological Field Station in southern Costa Rica at elevations between 1025 and 1200 m. This site is dominated by a mix of primary and secondary wet forest with a canopy up to 30 m tall and a midcanopy layer comprised of broadleaf trees, palms, and tree-ferns. Annual precipitation ranges from 3500-4000 mm and mean annual temperature at Las Cruces Biological Station is ∼21°C [24]. The bird checklist maintained by OTS for the station is comprised of over 400 species [25]. The Hawai’i study site was within the Maulua tract of Hakalau Forest National Wildlife Refuge on the island of Hawai’i. The canopy at this site is dominated by a *Metrosideros polymorpha*-*Acacia koa* trees up to 25 m tall with a mid-canopy of at least six native tree and tree-fern species. Mean annual precipitation is approximately 2500 mm and mean annual daily temperature is approximately is ∼15 °C at the study site [26]. Hakalau contains the most intact forest bird community remaining in the state with nine native species, plus an additional four non-native species that are common in the forest. Within each site, we recorded bird songs using automated acoustic recorders (SM2 Wildlife Acoustics Inc.) placed ∼1 meter above the ground at six different locations separated by at least 200 m. Recordings were made for 3-5 days at each location, depending on weather conditions. We programmed the acoustic recorders to record at 5-minute intervals (5-minutes on and 5-minutes off) throughout the day during the breeding season months of June through July 2012 in Costa Rica and March through April 2015 in Hawai’i. Recordings were made in WAV file format at a sampling rate of 44.1 kHz using a single omnidirectional microphone (SMX-II Wildlife Acoustics) with a sensitivity of −35 dBV/Pa and frequency response of 20-20,000 Hz. This study relied solely on the use of passive acoustic monitors placed in the forest to collect acoustic data. No animals were captured, handled, housed, monitored, or followed. No University IACUC permits were required for this study. Permission to conduct field work was given by landowners in both Costa Rica (Organization for Tropical Studies) and Hawai’i (Hakalau Forest National Wildlife Refuge).

### Acoustic and statistical analysis

All recordings were visualized on a spectrogram using Raven Pro 1.5 software (Bioacoustics Research Program 2017). Within each forest, we randomly selected one day at each of the six recording locations. We then selected three 5-minute recordings per day during the dawn chorus that did not contain rain or overlap with the chorus of cicada species such as *Zammara smaragdina* which can alter the timing and frequency of bird vocalizations in Costa Rica [22]. Interference by cicada choruses resulted in two sites in Costa Rica with only two 5-min recordings available.

We identified the bird species associated with each vocalization, then used the selection tool in RavenPro 1.5 to determine the minimum and maximum frequency and signal start and end time for all vocalizations within each 5-min recording period. Only selections > 5dB of the background noise of each recording were included in the analysis. All signals were measured using a Hann window type and a window size of 23.2 ms, window overlap of 50%, and DFT (discreet Fourier transform) size of 1024 samples [27]. Because signal characteristics can vary somewhat as a function of distance from the sender to the recorder, we calculated a mean minimum and maximum frequency for the signal of each species, and used this to calculate each signal’s average frequency range. We then created a frequency accumulation curve with frequency on the x-axis and the number of species using at least a portion of each frequency bin of 100 Hz on the y-axis. For each of the six recording locations, we calculated total species richness across the 5-minute time periods, and vocalization rate as the total number of vocalizations within a 5-min time period. We examined differences in species richness and vocalization rate between Hawai’i and Costa Rica using general linear mixed models and a Poisson link function in R (R Core Development Team 2018, version 3.5.1) with “forest” (Costa Rica, Hawai’i) as a fixed variable and “recorder site” as a random variable.

We used a null model approach (e.g., Masco et al. [28]) to test the ANH. For each 5-min recording period, we computed (i) the temporal and frequency overlap between each possible pair of vocalizations, and (ii) the total possible number of vocalization pairs as *n!/(2!(n-2)!*, where n = the number of vocalizations in a 5-min period (e.g., if n = 400, then the number of possible pairs is 79800). For each forest (Costa Rica, Hawai’i), the observed proportion of overlapping vocalization was computed as the sum of the number of overlapping pairs of vocalizations (i.e., pair of vocalizations with a spectral and temporal overlap > 0) divided by the sum of the total possible number of vocalization pairs. These observed proportions of overlapping vocalizations were then compared with the proportions obtained after randomizing the beginning of each vocalization 500 times but keeping the signal duration and average frequency range unchanged. Significantly fewer observed vs. expected overlaps between signals was considered evidence that birds partition acoustic space in each community. These analyses were performed separately for conspecific and heterospecific pairs of vocalizations. For Hawaii, we also examined whether there were differences in observed vs. expected overlap between pairs of native species vocalizations, pairs of alien species vocalizations, and mixed pairs of vocalizations (between one alien and one native species). We verified the results from the null model approach described above by using an additional method in which we visually scanned a set of 1-second “soundslices” from each recording to determine the proportion of overlapping vs. non-overlapping signals (see Supplemental Materials S1).

## Results

We detected 10 vocalizing species on our recordings from Hawaii and 47 from Costa Rica, though a number of the vocalizations from Costa Rica, primarily short calls, could not be identified to the species level (Table 1). There was a total of 4485 vocalizations detected with a duration of 6098 s from Hawaii and 2880 vocalizations with a duration of 1613 s from Costa Rica (Table 1). Three of the 10 species from Hawai’i were non-native and accounted for < 10% of total vocalizations. Recording locations in Costa Rica had more than double the median number of species (median = 17.5, range = 11 to 27) compared to locations in Hawai’i (median = 8, range = 5 to 8). We found a high degree of overlap in frequency use by species in both study areas with the most commonly used frequencies in Costa Rica ranging from 3000-3500 Hz (25 species) and from approximately 3000-6000 Hz (eight species) in Hawaii (Fig. 1). The mean (±SD) length of vocalizations was similar between study areas (1.13±0.96 seconds vs. 1.15±0.59 seconds in Costa Rica and Hawaii respectively). There was little difference in vocalization rate, indicating that the number of vocalizations per 5-minute period was similar between the two locations (z=1.77, P=0.077; Fig. 2A). As expected, there were significantly more species vocalizing per 5-minute recording period in Costa Rica than in Hawaii (z=-3.185, P= 0.001; Fig. 2B).

**Table 1.** The identity, total number of vocalizations detected, and total duration (in seconds) of vocalizations detected within 5400 and 4800 seconds of recordings analyzed for Hawaii and Costa Rica respectively.

| Site | Common name | Scientific name | Total vocalizations | Total duration |
| --- | --- | --- | --- | --- |
| CR | Rufous-tailed hummingbird | *Amazilia tzacatl* | 107 | 42.15 |
| CR | Orange-billed sparrow | *Arremon aurantirostris* | 143 | 143.46 |
| CR | Costa Rican brush finch | *Arremon costaricensis* | 84 | 66.59 |
| CR | Orange-billed nightingale thrush | *Catharus aurantirostris* | 231 | 215.01 |
| CR | Dusky antbird | *Cercomacroides tyrannina* | 1 | 1.77 |
| CR | Golden-olive woodpecker | *Colaptes rubiginosus* | 22 | 50.68 |
| CR | White-ruffed manakin | *Corapipo altera* | 11 | 4.7 |
| CR | Red-faced spinetail | *Cranioleuca erythrops* | 3 | 6.18 |
| CR | Little tinamou | *Crypturellus soui* | 4 | 17.23 |
| CR | Yellow-bellied elaenia | *Elaenia flavogaster* | 13 | 6.53 |
| CR | Spot-crowned euphonia | *Euphonia imitans* | 7 | 1.97 |
| CR | Black-faced antthrush | *Formicarius analis* | 27 | 38.21 |
| CR | Ruddy quail-dove | *Geotrygon montana* | 2 | 3.62 |
| CR | Red-crowned ant-tanager | *Habia rubica* | 17 | 4.32 |
| CR | White-breasted wood wren | *Henicorhina leucosticta* | 1052 | 409.67 |
| CR | Lesser greenlet | *Hylophilus decurtatus* | 27 | 12.95 |
| CR | Grey-chested dove | *Leptotila cassini* | 75 | 94.87 |
| CR | Scale-crested pygmy tyrant | *Lophotriccus pileatus* | 111 | 75.08 |
| CR | White-whiskered puff bird | *Malacoptila panamensis* | 15 | 10.82 |
| CR | Greenish elaenia | *Myiopagis viridicata* | 1 | 0.97 |
| CR | Buff-rumped warbler | *Myiothlypis fulvicauda* | 7 | 18.19 |
| CR | Scaley-breasted hummingbird | *Phaeochroa cuvierii* | 28 | 13.74 |
| CR | Green hermit | *Phaethornis guy* | 19 | 2.59 |
| CR | Stripe-throated hermit | *Phaethornis striigularis* | 589 | 110.44 |
| CR | Rufous breasted wren | *Pheugopedius rutilus* | 29 | 59.23 |
| CR | Squirrel cuckoo | *Piaya cayana* | 3 | 2.91 |
| CR | Olivaceous piculet | *Picumnis olivaceus* | 1 | 1.39 |
| CR | Chestnut-backed antbird | *Poliocrania exsul* | 1 | 0.77 |
| CR | Chestnut-mandibled toucan | *Ramphastos ambiguous swainsonii* | 65 | 33.78 |
| CR | Long-billed gnatwren | *Ramphocaenus melanurus* | 12 | 23.52 |
| CR | Rufous mourner | *Rhytipterna holerythra* | 7 | 7.48 |
| CR | Tropical parula | *Setophaga pitiayumi* | 10 | 18.87 |
| CR | Russet antshrike | *Thamnistes anabatinus* | 59 | 21.3 |
| CR | Yellow-olive flycatcher | *Tolmomyias sulphurescens* | 9 | 6.37 |
| CR | White-throated thrush | *Turdus assimilis* | 23 | 45.7 |
| CR | Clay-colored thrush | *Turdus grayi* | 40 | 23.21 |
| CR | Paltry tyrannulet | *Zimmerius vilissimus* | 18 | 9.22 |
| HI | Northern cardinal* | *Cardinalis cardinalis* | 77 | 87.2 |
| HI | Hawai'i 'Elepaio | *Chasiempis sandwichensis* | 39 | 12.19 |
| HI | Hawai'i 'Amakihi | *Chlorodrepanis virens* | 359 | 492.35 |
| HI | 'I'iwi | *Drepanis coccinea* | 2154 | 3221.59 |
| HI | 'Apapane | *Himatione sanguinea* | 800 | 1357.61 |
| HI | Red-billed leiothrix* | *Leiothrix lutea* | 65 | 99.29 |
| HI | Hawai'i 'Ākepa | *Loxops coccineus* | 7 | 3.4 |
| HI | 'Alawī | *Loxops mana* | 48 | 84 |
| HI | 'Ōma'o | *Myadestes obscurus* | 737 | 380.45 |
| HI | Warbling white-eye* | *Zosterops japonicus* | 203 | 358.49 |
| CR | Unknown 9 |  | 2 | 1.72 |
| CR | Unknown 14 |  | 6 | 1.25 |

**Fig 1.**
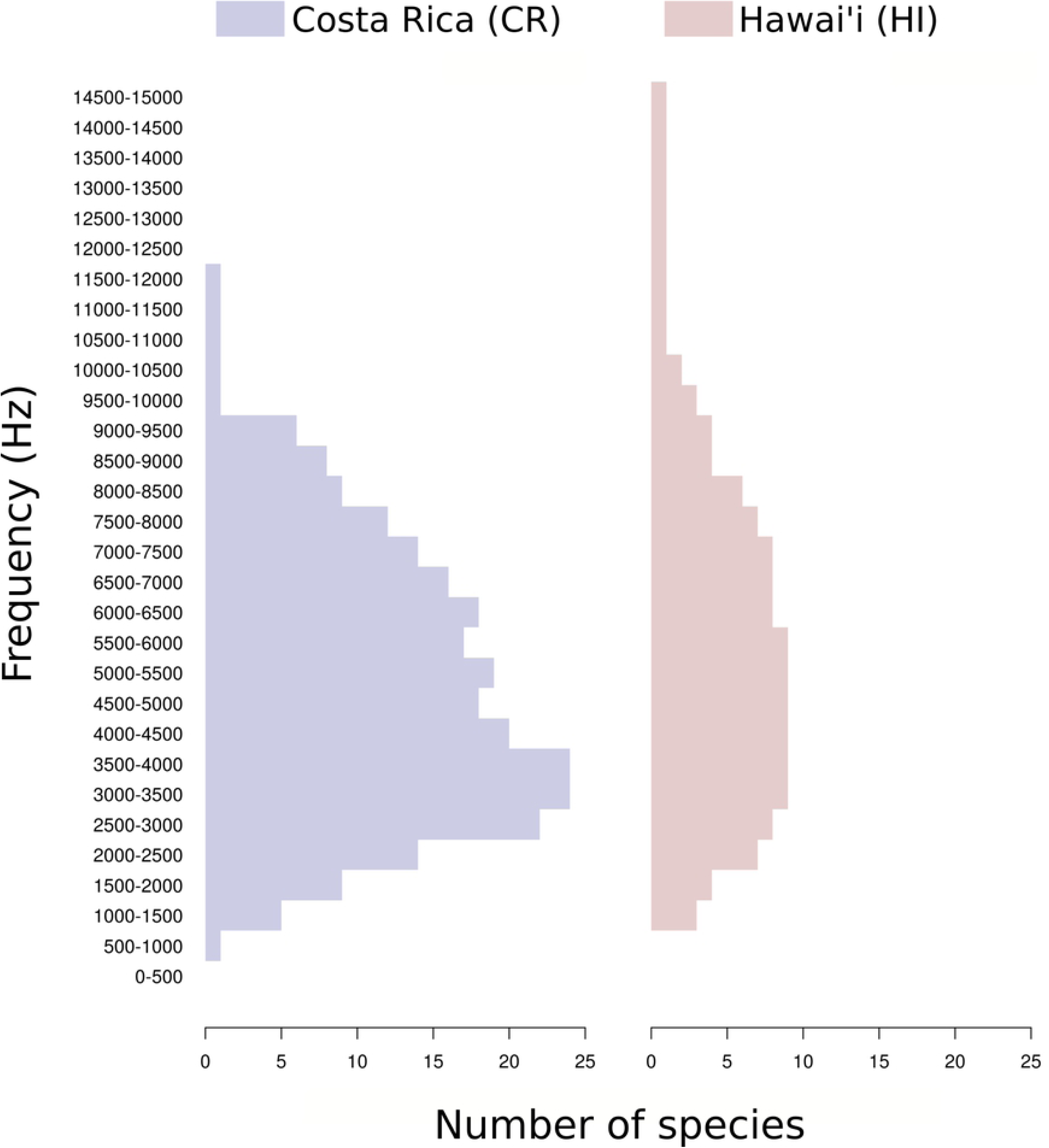
Accumulated frequency use based on mean minimum and maximum frequencies of vocalizations for 47 species in Costa Rica and 10 species in Hawaii.

**Fig 2.**
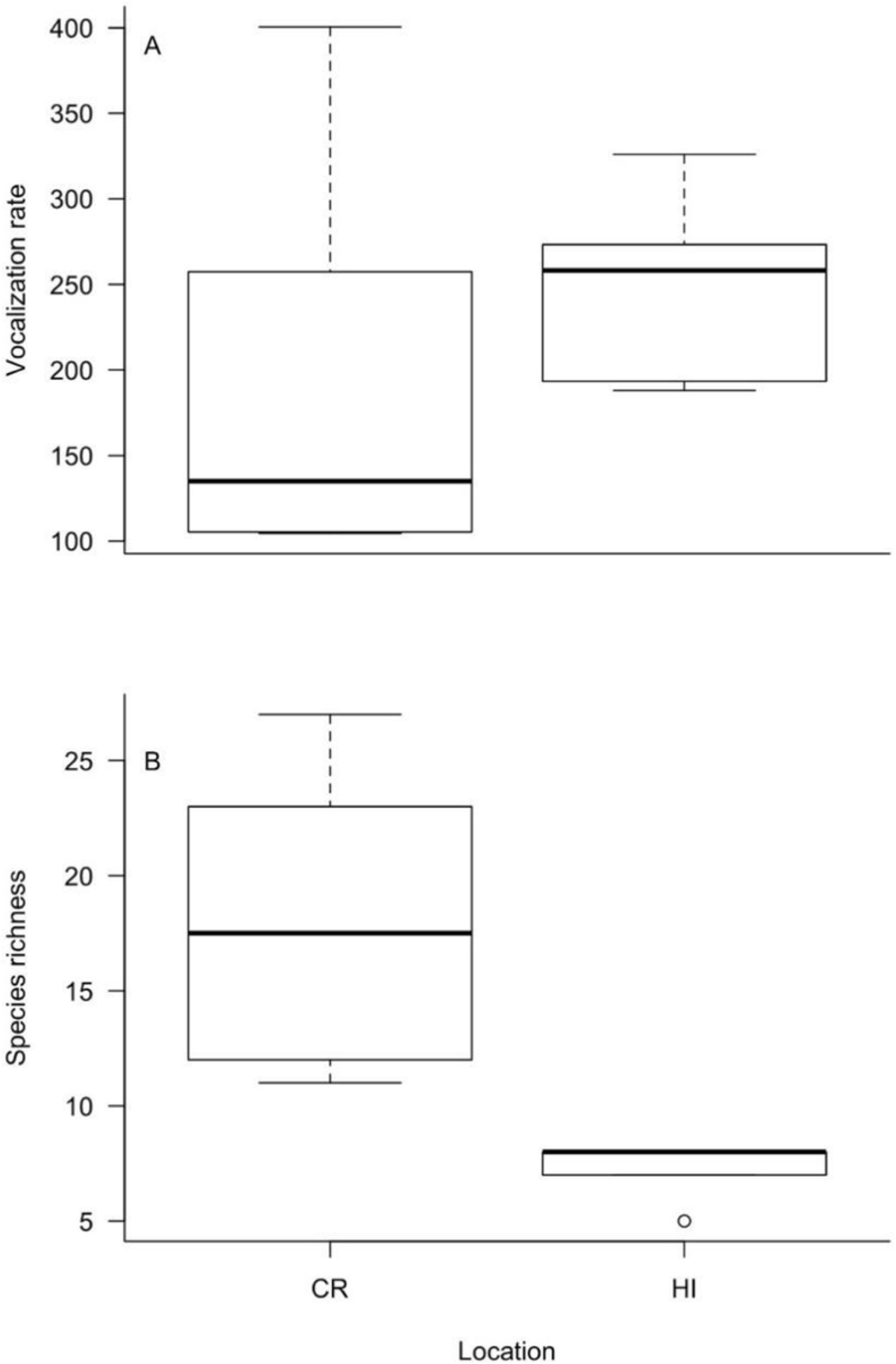
Boxplots of the difference in A) vocalization rate (vocalizations per 5-minutes) and B) species richness between the tropical forests in Costa Rica and Hawai’i. The whiskers are representative of the 10^th^ and 90^th^ percentiles, the boxes show the 25^th^ and 75^th^ percentiles, the mean value is indicated by the solid black line and circles represent outliers.

Observed intra- and interspecific signal overlap (in time and frequency combined) was far less than expected by chance in both Costa Rica and Hawaii (Fig. 3). Indeed, there were no randomly generated iterations of expected values that coincided with the number of observed signal overlaps. Observed signal overlap (time, time and frequency combined) was generally greater in Hawaii than in Costa Rica (Fig. 3). In particular, observed interspecific overlap in time and frequency combined was more than an order of magnitude greater in Hawaii (Fig. 3). Observed intraspecific overlap was rare in both study locations, but the difference between inter- and intraspecific overlap for time and frequency combined was much greater in Hawaii than Costa Rica. Finally, we did not detect a significant difference in the observed vs. expected level of overlap between native and non-native species in Hawaii (Fig. 4).

**Fig 3.**
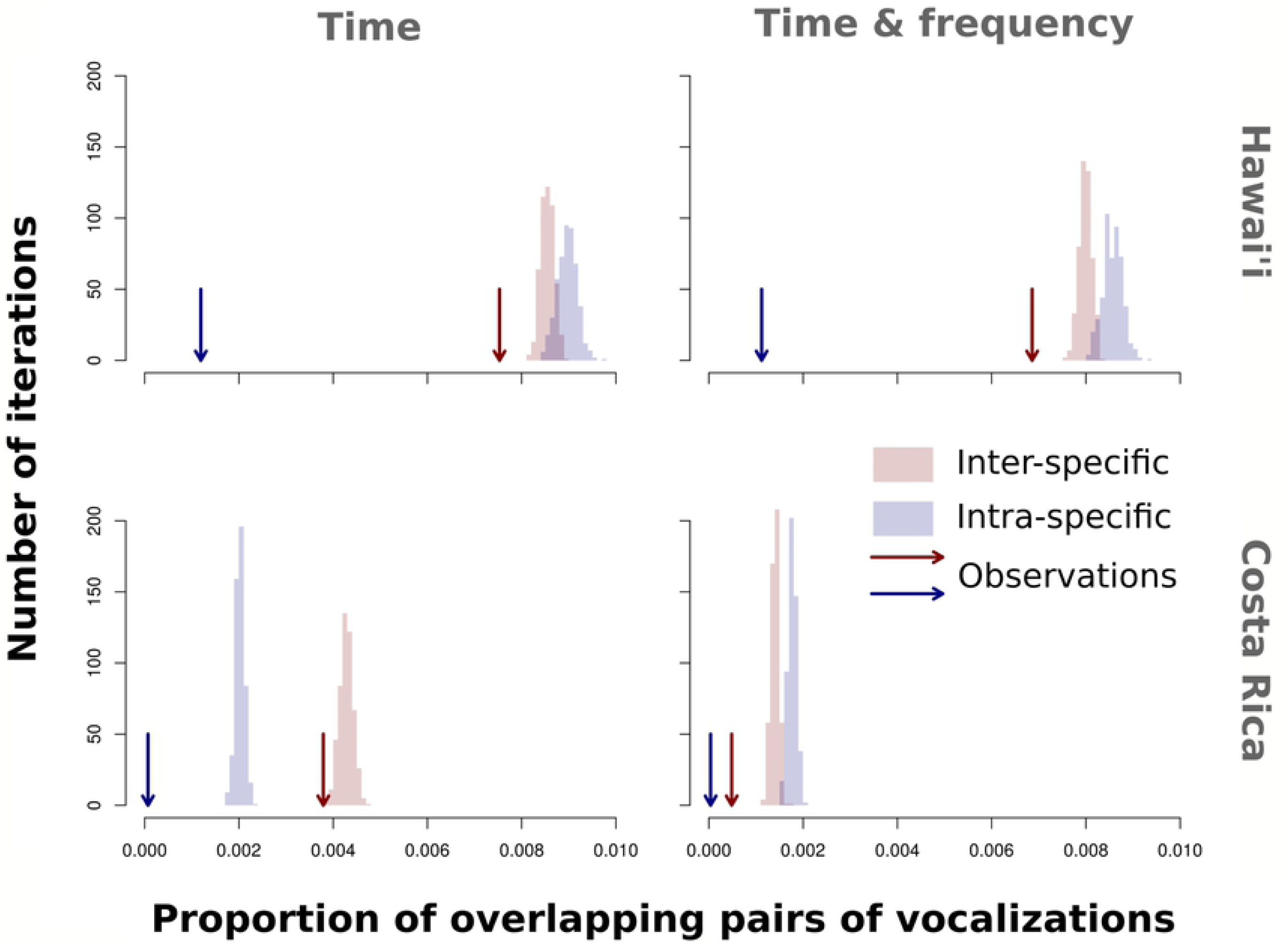
Comparison between the observed proportion of overlapping pairs of vocalizations (arrows) and the expected proportion of overlapping pairs of vocalizations under null models (bars, 500 iterations) for both interspecific and intraspecific vocalization pairs in Hawai’i and Costa Rica.

**Fig 4.**
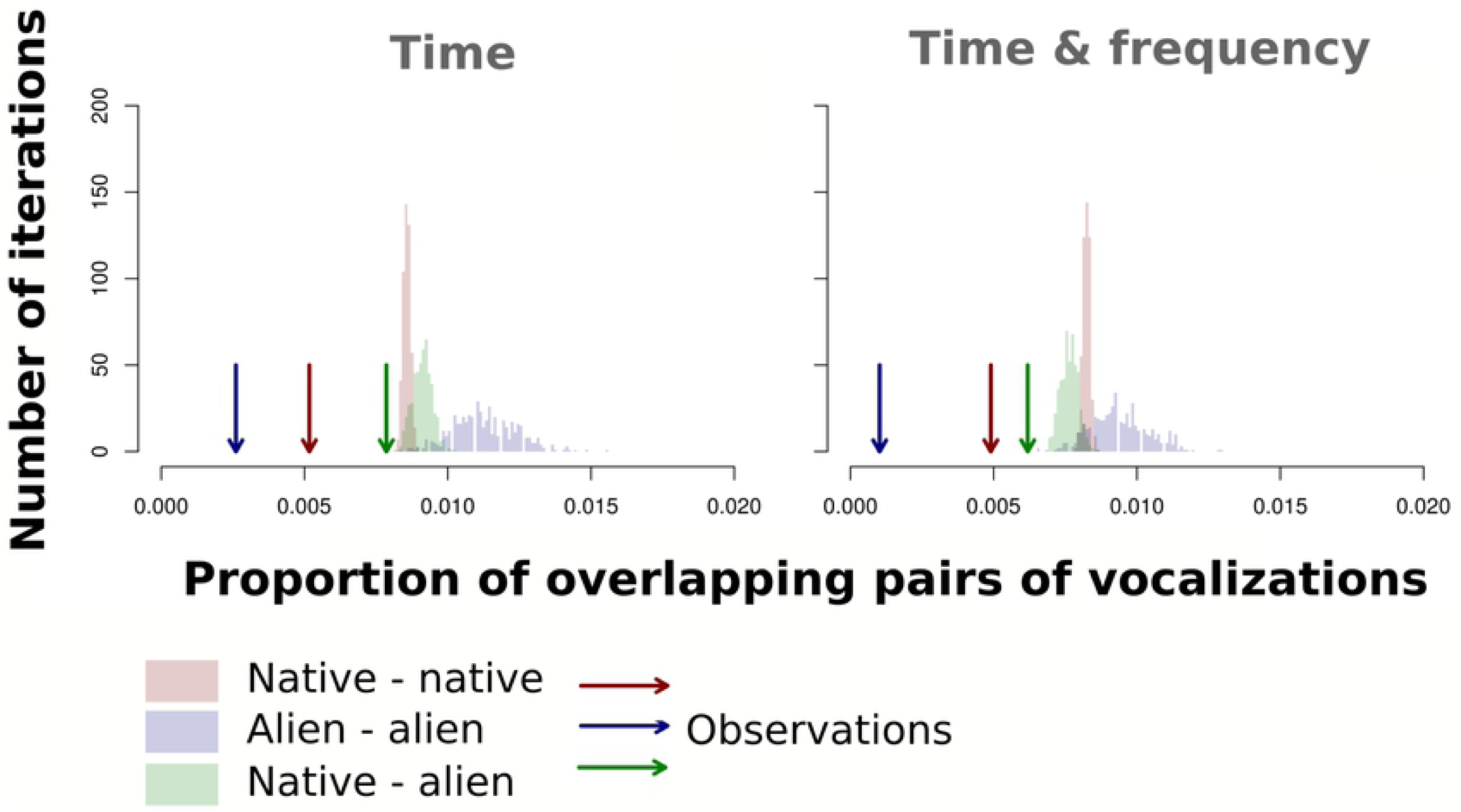
Comparison between the observed proportion of overlapping pairs of vocalizations (arrows) and the expected proportion of overlapping pairs of vocalizations under null models (bars, 500 iterations) among native and non-native (alien) vocalization pairs in Hawai’i.

## Discussion

As predicted under ANH, we found significantly less temporal and spectral overlap of acoustic signals than expected by chance in wet forest bird communities in Costa Rica and Hawaii. These findings constitute strong support that there is competition for acoustic space in species-rich signaling communities because they are based on null models generated at the level of each recording period. Past studies using null models have found support for spectral partitioning in anuran [11], orthopteran [10], and even spectral [8] or temporal [29] clustering of acoustic signals at the community level. However, evidence for simultaneous spectral and temporal partitioning at the community level, critically important to demonstrating acoustic niche partitioning in real time, has been lacking. It was also notable that observed partitioning was more pronounced within than among species in both locations (Fig. 3). This indicates that there may be greater selection pressure to reduce overlap with conspecifics, possibly because intraspecific signals are far more similar than interspecific ones.

While we found evidence for acoustic niche partitioning in both communities, and despite similarities in signaling rate and in the range of spectral bandwidths used, we found strong differences between communities in the use of acoustic space. Overlap between different signals was very rare in Costa Rica but occurred within approximately 37 % of the sound slices examined in Hawaii (Supplementary Material S1). These findings could be strengthened by increasing replication across both regions, but they indicate that greater species richness in Costa Rica may have led to stronger selection for reducing interspecific signal overlap there. In addition, the island of Hawaii is geologically younger than much of Costa Rica (∼0.5 vs 2.5-15 my; [30]) so there are likely differences in length of time available for the evolution of signal character displacement between locations. There may be a phylogenetic effect as well – the Costa Rican bird community represents a far greater diversity of avian lineages than Hawaii, where most of the current species have evolved through adaptive radiation from just a handful of colonizations from distant continents [31]. Differences in signal function may also partially explain the different levels of signal overlap between the two communities. Most of the Costa Rica species sampled here maintain year-round territories [32], whereas most of the Hawaiian species are generally non-territorial outside the breeding season. The function of song may be more associated with promoting social cohesion, with less selection against reducing interspecific signal overlap, in communities with higher densities of individual species that are non-territorial.

We focused our analyses on the period during which birds most actively vocalize in each forest. This period differs between Hawaii and Costa Rica, possibly as a function of the presence of other calling taxa. In Costa Rica, the night-time soundscape is dominated by arthropods and anurans, with relatively little bird activity. The dawn chorus for birds begins soon after first light, and continues for approximately 1.5-3 hours, when the daily chorus of cicadas begins. Past work has shown that cicada choruses dominate the frequency bands commonly used by birds and may effectively shut down communication for many bird species in Costa Rica [22]. The dawn period is when birds have the least apparent competition with other calling organisms and the avian community may partition acoustic space to a greater degree than any other time period during the day. Conversely, Hawaii has no native anurans, no cicadas, and far fewer arthropods that signal acoustically during the day. Perhaps for this reason, the morning chorus period may last for up to five hours or more in forests dominated by native Hawaiian bird species such as Hakalau Forest NWR.

An important prediction that is based on the Acoustic Niche Hypothesis is that habitats with less altered species assemblages will exhibit greater levels of acoustic partitioning than habitats with more altered species assemblages [6]. For example, vocalizations of the non-native *Leiothrix lutea* frequently overlap with those of native species, and thus reduce acoustic partitioning at the community level in an Italian shrubland [33]. Most bird communities in Hawaii are comprised of a mix of native species and non-native species that have been introduced within the past 100 years. In our study, 30% (3 out of 10) of the bird recorded in Hawaii were non-native species (including *Leiothrix lutea*), whereas none in Costa Rica are considered non-native. We did not detect a significant difference in the observed vs. expected level of overlap between native-native vs. native-non-native species (Fig. 4), however Hakalau Forest is notable for its lack of non-native bird species relative to most other areas in Hawaii and their vocalizations comprised less than 10% of the total. Future studies could compare overlap between native and non-native dominated bird communities in Hawaii as a test of this prediction of ANH.

## Conclusions

Our finding that birds simultaneously partition acoustic space both spectrally and temporally, and that the strength of this partitioning varies between two communities, adds to the growing body of knowledge that this may be a general phenomenon in other areas with high densities of signaling individuals. The null model approach we used here would likely be a useful method for testing the acoustic niche hypothesis in these areas.

## Acknowledgements

We thank Kary Perez and Michael Atencio-Picado for assistance with acoustic analysis. Also, Zak Zahawi and the staff at the Organization for Tropical Studies Las Cruces Field Station for facilitating field work in Costa Rica, and Steve Kendall of the US Fish and Wildlife Service for permits and access to field sites in Hawaii.

